# Structural basis for the inhibition of the papain-like protease of SARS-CoV-2 by small molecules

**DOI:** 10.1101/2020.07.17.208959

**Authors:** Ziyang Fu, Bin Huang, Jinle Tang, Shuyan Liu, Ming Liu, Yuxin Ye, Zhihong Liu, Yuxian Xiong, Dan Cao, Jihui Li, Xiaogang Niu, Huan Zhou, Yong Juan Zhao, Guoliang Zhang, Hao Huang

## Abstract

SARS-CoV-2 is the pathogen responsible for the COVID-19 pandemic. The SARS-CoV-2 papain-like cysteine protease has been implicated in virus maturation, dysregulation of host inflammation and antiviral immune responses. We showed that PLpro preferably cleaves the K48-ubiquitin linkage while also being capable of cleaving ISG15 modification. The multiple functions of PLpro render it a promising drug target. Therefore, we screened an FDA-approved drug library and also examined available inhibitors against PLpro. Inhibitor GRL0617 showed a promising IC_50_ of 2.1 μM. The co-crystal structure of SARS-CoV-2 PLpro-C111S in complex with GRL0617 suggests that GRL0617 is a non-covalent inhibitor. NMR data indicate that GRL0617 blocks the binding of ISG15 to PLpro. The antiviral activity of GRL0617 reveal that PLpro is a promising drug target for therapeutically treating COVID-19.

**One Sentence Summary:** Co-crystal structure of PLpro in complex with GRL0617 reveals the druggability of PLpro for SARS-CoV-2 treatment.

---

The COVID-19 pandemic has caused devastating damage to the world and it has resulted in over 12 million confirmed cases and over half a million death as of July 14, 2020 (*1*). The novel SARS-CoV-2 coronavirus is the etiological agent responsible for the pandemic, and it belongs to the beta coronavirus family (*2-4*). Similar to the other two known beta coronaviruses, i.e. SARS and MERS, SARS-CoV-2 also causes severe acute respiratory syndromes (*5, 6*). Unexpectedly, SARS-CoV-2 has been reported to have more mild symptoms but much higher transmission rate (*7, 8*), therefore it has caused the biggest catastrophe to the world healthcare since the Spanish flu in 1918-1920 (*9*). There are currently no FDA-approved drugs for SARS, MERS or SARS-CoV-2. Remdesivir, an anti-HIV drug developed by Gilead, was given Emergency Use Authorization (EUA) permission by the U.S. Food and Drug Administration (FDA) for COVID-19 treatment (*10*). Meanwhile, the discovery of antibodies or vaccines for COVID-19 is still in progress. Therefore, anti-SARS-CoV-2 drugs are urgently needed.

As a positive strand RNA Virus, SARS-CoV-2 encodes two functional proteases, i.e. the papain-like protease (PLpro) and the 3-chymotrypsin-like cysteine protease (Mpro or 3CLpro). The major function of Mpro is to cleave the viral polyprotein, which is critical for virus maturation, replication and invasion. Several potent Mpro covalent inhibitors have been reported and their co-crystal structures provided opportunities for structure-based drug optimization (*11-15*). SARS-CoV PLpro is a cysteine protease with multiple major functions, including processing of the viral polyprotein chain for viral protein maturation, dysregulating host inflammation responses through deubiquination and impairing the host type I interferon antiviral immune responses by removing ISG15 modifications from host proteins (*16-18*). SARS-CoV-2 PLpro (MW = 35.6 kDa) contains two domains including an N-terminal Ubiquitin-like domain and a C-terminal ubiquitin specific protease (USP) domain with implicated catalytic functions of cleaving ubiquitin (Ub) or ISG15 modifications from host proteins (*19*).

Besides Mpro, PLpro has been considered another promising target for drug discovery to treat COVID-19 (*20*). Due to the urgent situation of the pandemic, the strategy of repurposing approved drugs or optimizing new compounds has been employed to fight COVID-19 (*15, 21, 22*).

## Results

SARS-COV-2 PLpro and SARS PLpro are closely related homologs sharing ∼83% sequence identity. It has been reported that SARS-CoV PLpro preferably cleaves K48 linkages of polyubiquitin chains as well as ISG15 modifications from host proteins (*23, 24*). Using SDS-PAGE and silver-staining, we investigated the cleavage preference of SARS-COV-2 PLpro using *in vitro* deubiquitination assays for di-Ub and tetra-Ub samples (Fig. 1). As shown in Fig. 1A, di-Ub of various linkage types including K6, K11, K27, K29, K33, K48 and K63 were treated with SARS-COV-2 PLpro. Within 30 min, nearly 50% K48-di-Ub was cleaved into mono-Ub, while K63- and K11-diUb were also cleaved but to a much lesser extent. The other four linkages, namely K6, K27, K29 and K33 were uncleaved. Apparently, K48 was the preferred linkage for SARS-COV-2 PLpro. We further tested the length preference of the polyUb chain for SARS-COV-2 PLpro. For the K48 linkage, PLpro showed a faster cleavage rate for tetra-Ub with over than 50% tetra-Ub cleaved at 15 min, while it took over 30 minutes for di-Ub to be cleaved over 50% (Fig. 1B). For K63, neither di-Ub nor tetra-Ub were significantly cleaved within one hour. Apparently, the K63 linkage was less favored by SARS-COV-2 PLpro compared with K48 for either di-Ub or the longer tetra-Ub (Fig. 1B and C). Western blot experiments of the PLpro cleavage of K11-, K63- and K48-tetra-Ub also showed that K48 was more favored than K63, and that the K11 and M1 linkage could not be cleaved by PLpro within 30 min. Under short exposure time of 0.5 sec, cleaved tri-Ub and di-Ub were observed only for K48 (Fig. 1D) and the mono-Ub was observed for K48 after an exposure time of 2 sec (Fig. S1).

**Figure 1.**
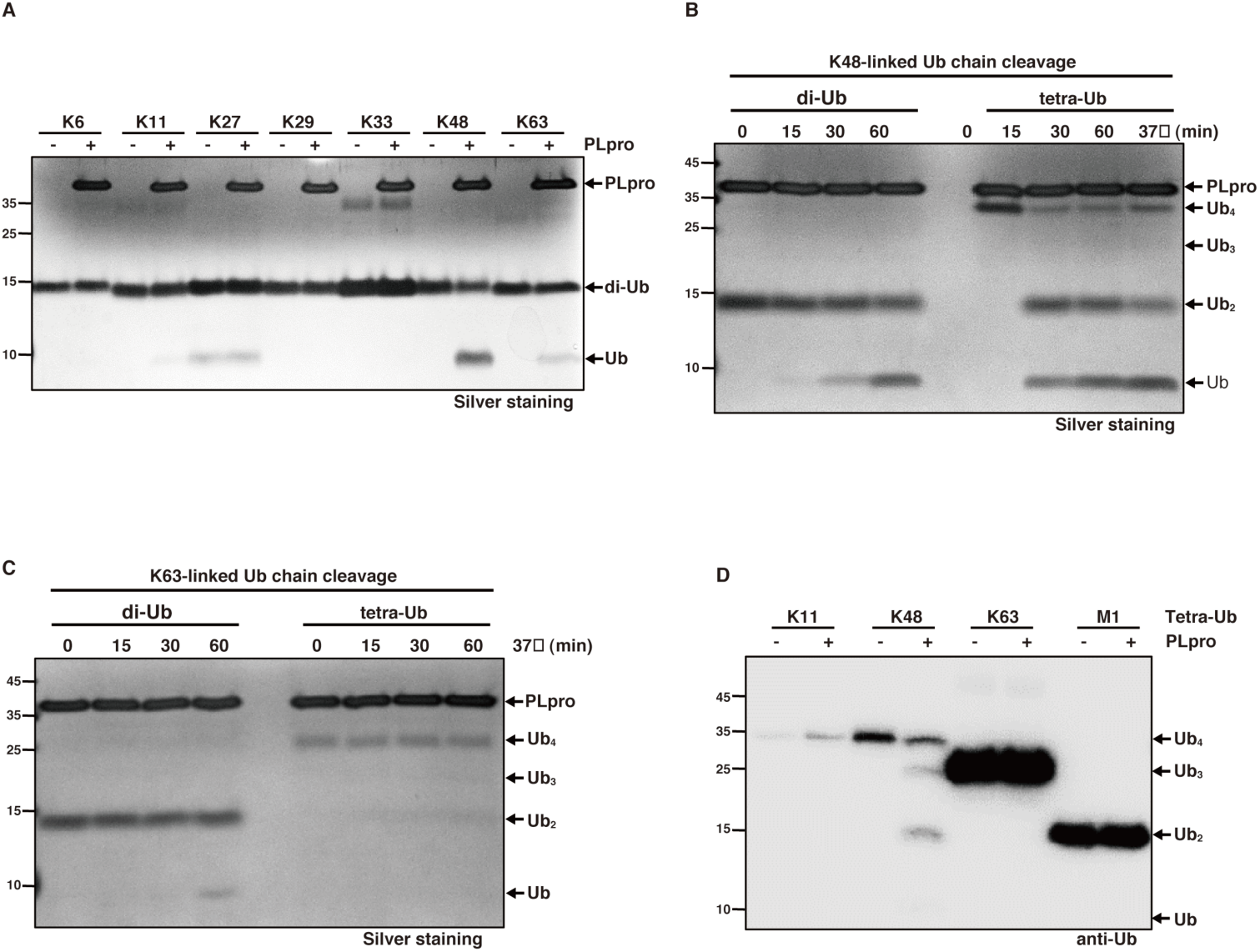
SARS-COV-2 PLpro prefers K48 over other di-Ub linkages and K48 tetra- Ub was more favored than di-Ub. (A) Seven linkage types (K6, K11, K27, K29, K33, K48 and K63) of di-Ub were incubated with purified recombinant PLpro (10 nM) for 30 min at 37°C, then analyzed by SDS-PAGE and silver staining. (B-C) K48- and K63-di-Ub or tetra-Ub (1 mM) were incubated with purified recombinant PLpro (10 nM) for indicated time, then analyzed by SDS-PAGE and silver staining. (D) PLpro was incubated with K11-, K48-, K63-tetra-Ub and M1 di-Ub for 30 min at 37°C, then analyzed by SDS-PAGE and immunoblotting with 0.5 sec exposure.

To validate that PLpro is a potential drug target and to repurpose existing drugs to treat COVID-19, we initiated screening of a 2111-compound FDA approved drug library against SARS-COV-2 PLpro. First, we set up an FRET assay to characterize the enzymatic activity of SARS-COV-2 PLpro based on an established assay for SARS-Cov PLpro (*25*). The recombinant full-length SARS-COV-2 PLpro protein was expressed in *E*. *coli* and subsequently purified. A commercially available fluorogenic peptide substrate RLRGG-AMC, representing the C-terminal residues of ubiquitin, was used to report the enzymatic activity of PLpro. The 1^st^ round of screening provided ∼30 compounds with over 50% inhibition at 100 μM. Hits from the 1^st^ round of screening went into the 2^nd^ round of validation using the same enzymatic assay. After removing compounds with poor solubility, strong reactivity and low intrinsic fluorescence, seven relatively potent compounds were measured for IC_50_ (n=3). These 7 drugs showed modest IC_50_ values ranging from 29 to 91 μM (Fig. S3). Although these compounds can potentially provide a starting point for further optimization, their low potency imply a need of large amounts of resources and time input. Therefore, we cherry-picked GRL0617 from promising SARS-Cov PLpro inhibitors based on the structural similarity between the SARS-CoV and SARS-CoV-2 PLpro proteins (*25*). The *in-vitro* IC_50_ of GRL0617 against SARS-COV-2 PLpro was 2.1 ± 0.2 μM, suggesting a promising lead compound and therefore it was subjected to further structural and mechanistic studies (Fig. 2A).

**Figure 2.**
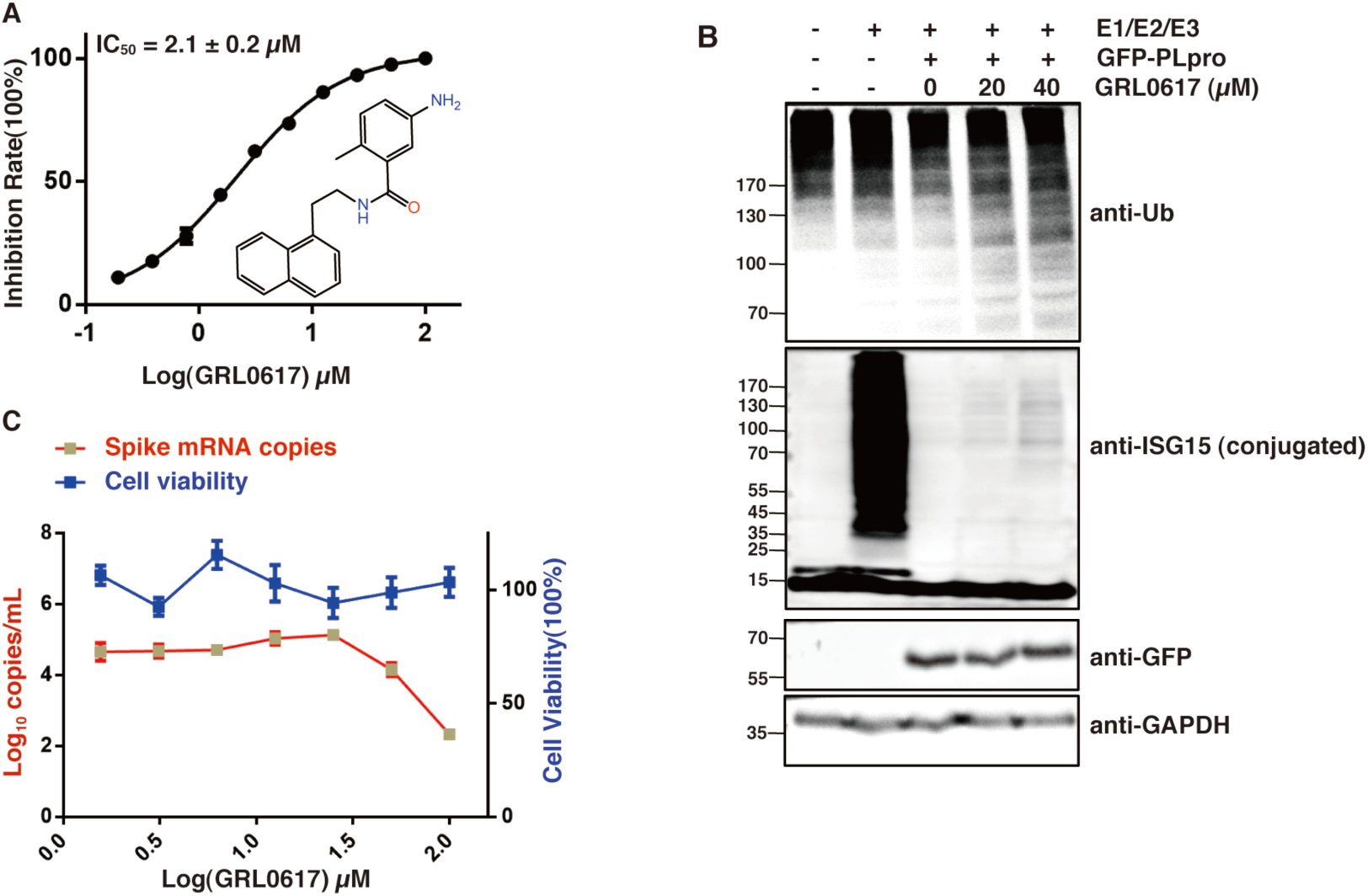
Inhibitory activity of GRL0617 against SARS-COV-2 PLpro. (A) The inhibitory activity of GRL0617 against PLpro was measured using peptide substrate RLRGG-AMC as substrate. IC_50_ was presented as mean ± SEM (n=3). (B) HEK293T cells were transfected for 24?h with plasmids encoding GFP-PLpro, ISG15, E1(Ube1L)/E2(UbcH8)/E3(HECR5) enzymes, alone or in combination. Cells were treated for an additional 24?h with indicated concentrations of GRL0617. Cell lysates were subjected to immunoblotting with anti-ubiquitin, anti-ISG15, anti-GFP antibodies. GAPDH served as a loading control. (C) Antiviral activity of GRL0617 on SARS-CoV-2 and the cytotoxicity of GRL0617 on Vero E6 cells. Vero E6 cells were infected with SARS-CoV-2 using a multiplicity of infection (MOI) of 0.01. The quantification of absolute viral RNA copies (per mL) in the supernatant at 48 h post infection was determined by qRT-PCR analysis. The cytotoxicity of GRL0617 on Vero E6 cells were measured using CCK8. All data are shown as mean± SEM, n = 3.

To assess whether GRL0617 can inhibit the *in vivo* deubiquitinating and deISGylating activity of SARS-COV-2 PLpro, we transfected HEK293T cells with expression constructs of PLpro and the ISGylation machinery (Ube1L, UbcH8, HECR5 and ISG15) and treated with GRL0617 in a dose-depedent manner for 24 hours. Our data showed that SARS-COV-2 PLpro is capable of reverse polyubiquitination and ISGylation of cellular substrates (Fig. 2B). Moreover, GRL0617 clearly showed inhibition of SARS PLpro and resulted in recovered poly-ubiquitin-conjugates and ISG15-conjugates (Fig. 2B).

To further validate the potential of PLpro as an antiviral drug target, we tested GRL0617 for its inhibitory activity in Vero E6 cells infected with SARS-CoV-2 at a multiplicity-of-infection (MOI) of 0.01. The copy numbers of the viral spike protein mRNA was monitored to evaluate antiviral activity of the compound. Based on the dose-dependent response, GRL0617 showed a clear inhibition of viral replication and 100 μM of GRL0617 resulted in over 50% inhibition. No apparent cytotoxicity on Vero E6 cells was observed in our assay with concentrations up to 100 μM (Fig. 2C).

A co-crystal structure would be crucial to understand the mechanism of inhibition of SARS-COV-2 by GRL0617, therefore we determined the X-ray co-crystal structure of SARS-COV-2 PLpro C111S at 3.2 Å (Fig. 3A-E and Table S1). The crystal of PLpro/GRL0617 belongs to the space group I4122 with just one protein molecule in each asymmetric unit. GRL0617 resides in the S3 and S4 subsites of PLpro and is apart from the catalytical triad with a minimum distance of 10.5 Å to S111 in the C111S structure. Based on the binding site of GRL0617, it inhibits SARS-COV-2 PLpro in a non-covalent manner. The overall GRL0617 bound PLpro structure is very similar to the available apo-structure of C111S (PDB 6WRH) with a backbone RMSD of 0.76 Å, except for two residues on the BL2 loop, i.e. Y268 and Q269 (Fig. 3D). Upon binding GRL0617, the sidechain of Y268 and Q269 shifted towards GRL0617 to form polar and hydrophobic interactions with the compound and stabilize its binding (Fig. 3D). Specifically, the sidechain oxygen of Y268 forms a hydrogen bond with the NH_2_ group on the benzene ring of GRL0617, and the backbone NH of Q269 with the carbonyl oxygen of GRL0617 (Fig. 3B and C). In comparison with the apo-wild type (wt-, PDB 6W9C) or C111S (PDB 6WRH) structures of SARS-COV-2 PLpro, the BL2 loop of GRL0617 bound C111S structure shifted towards the compound and form a deeper pocket to better accommodate the compound (Fig. 3E and Fig. S4 A, D and E). Other polar interactions include the sidechain oxygen of Y264 and backbone carbonyl oxygen with the amide NH of GRL0617. In addition, hydrophobic integration also plays pivotal roles in GRL0617’s binding to PLpro. The naphthalene group of GRL0617 is buried in the pocket formed by aromatic residues Y264, Y268 and the hydrophobic sidechains of P247 and P248 (Fig. 3B and C).

**Figure 3.**
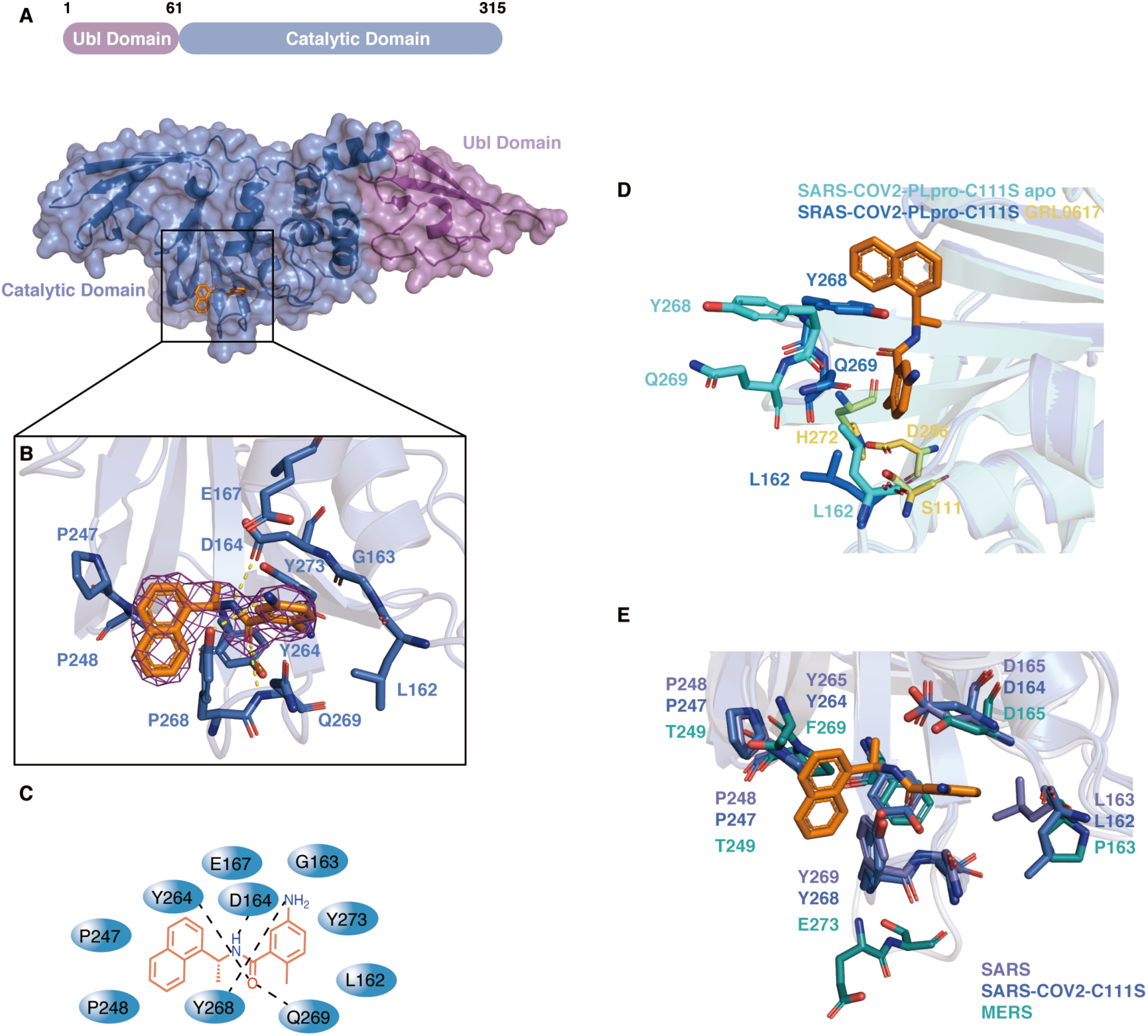
Structural and mechanistic analysis of SARS-COV-2 PLpro C111S in complex with GRL0617, and in comparison with SARS and MERS PLpro. (A) Surface and cartoon structures of SARS-CoV-2 PLpro in complex with GRL0617 (orange sticks) showing the N-terminal UBL domain (magenta) and C-terminal USP domain (marine). (B) The binding pocket of GRL0617 in PLpro. The PLpro residues involved in GRL0617 binding are shown as marine sticks. The 2Fo-Fc omit map (contour level = 1.6 σ, shown as purple mesh). (C) Schematic diagram of SARS-COV-2 PLpro C11S/GRL0617 interactions shown in (B). (D) Comparison of GRL0617-bound (blue) PLpro C111S and unbound (cyan) C111S (PDB ID: 6WRH) structures. Y268 and Q269 on the BL2 loop shifted towards GRL0617 upon binding. GRL0617 shown as orange sticks; The catalytic triad residues (S111 in place of C111, H272 and D286) are shown in yellow; Y268 and Q269 are shown in marine in bound state, and in cyan in unbound state. (E) Comparisons of the binding sites of SARS-CoV PLpro/GRL0617 (slate sticks, PDB ID: 3E9S)(*25*), SARS-COV2 PLpro C111S/GRL0617 (marine sticks) and MERS-CoV PLpro (deep teal sticks, PDB ID: 4RNA)(*29*).

Since GRL0617 is capable of inhibiting both SARS-CoV and SARS-CoV-2, it is of interest to see if it can be a broad spectrum antiviral inhibitor against other coronaviral PLpro proteins. It has been reported that naphthalene-based compounds have low to zero potency towards MERS PLpro(*26*). The superposition of the GRL0617/SARS-COV-2 PLpro and MERS PLpro indicated that the original pocket in MERS PLpro might be too shallow to allow GRL0617 to bind with extensive contacts, and the naphthalene moiety of GRL0617 would also be in steric clash with the pocket of MERS PLpro (Fig. S4C). In contrast to SARS and SARS-CoV-2, the BL-2 loop of MERS is one residue longer but it lacks the critical Y268 of SARS-COV-2 (Fig. S2). The extra residue of MERS PLpro may rearrange the hydrogen-bonding interaction network of the BL2 loop and the lack of the aromatic tyrosine clearly resulted in the removal of T-shaped Pi-Pi stacking and van der Waals interactions with the naphthalene group of GRL0617 (Fig. 3E and Fig.S4 A-F).

Because GRL0617 is a non-covalent inhibitor binding in the S3/S4 site of SARS-COV-2 PLpro, it is expected that GRL0617 would disrupt the interactions between ISG15 or Ub with PLpro. Solution state NMR was employed to characterize the binding of ISG15 or Ub to PLpro as well as the perturbation of their bindings by GRL0617. The 2-D NMR ^1^H,^15^N-HSQC spectrum of ^15^N-ISG15 (0.1 mM) shows typical features for a well folded protein with dispersed cross peaks (Fig. 4A). The addition of 0.15 mM SARS-COV-2 PLpro into 0.1 mM ^15^N-ISG15 caused drastic peak broadening and peak intensity loss, which is a characteristic of protein-protein interactions in the intermediate chemical exchange regime (Fig. 4B). Increasing concentrations of GRL0617 were added into the mix of 0.1 mM ^15^N-ISG15 and 0.15 mM of SARS-COV-2-PLpro, a dose dependent response of peak intensity recovery was evident, which indicated that GRL0617 competes with ISG15 for the binding site on PLpro, namely the S3 and S4 sites and blocked the binding of ISG15 to PLpro (Fig. S5A). The superposition of the HSQC spectra of 0.1 mM ISG15 only and the 0.1 mM/0.15 mM PLpro/0.25 mM GRL0617 mixture are essentially identical, which indicated that GRL0617 is a potent binder to PLpro and almost completely blocked the binding of ISG15 to PLpro at a molar ratio of 1.67 (0.25 mM /0.15 mM) (Fig. 4C and Fig. S5B). No peak shifting was observed in the superimposed HSQC (Fig. S5B), suggesting that GRL0617 is a *bona fide* PLpro binder because the HSQC spectrum of ^15^N-ISG15 is not disturbed at all by 2.5 excess molar ratio of GRL0617. In comparison with the complex structure of SARS-COV-2 PLpro with Ub (PDB ID: 6XAA) (Fig. 4F) or ISG15 (PDB ID: 6XA9) (Fig. 4E), GRL0617 blocked the access of the C-terminal tail of Ub and ISG15 to the catalytically active site of SARS-COV-2 PLpro. As suggested from the linkage cleaving preference experiments (Fig. 1), only K48 linkage can be effectively cleaved by SARS2-PLpro, therefore a very weak binding of monoUb to PLpro was expected. Indeed, titrations of PLpro into ^15^N-Ub caused very limited peak shifting even at molar ratio of 3, confirming the suspected weak binding (Fig. S6). Therefore, GRL0617 was not further titrated into the ^15^N-Ub/PLpro mixture. Taken together, our NMR and X-ray analysis indicates that GRL0617 is a potent PPI (protein-protein interaction) inhibitor for PLpro blocking the binding of ISG15.

**Figure 4.**
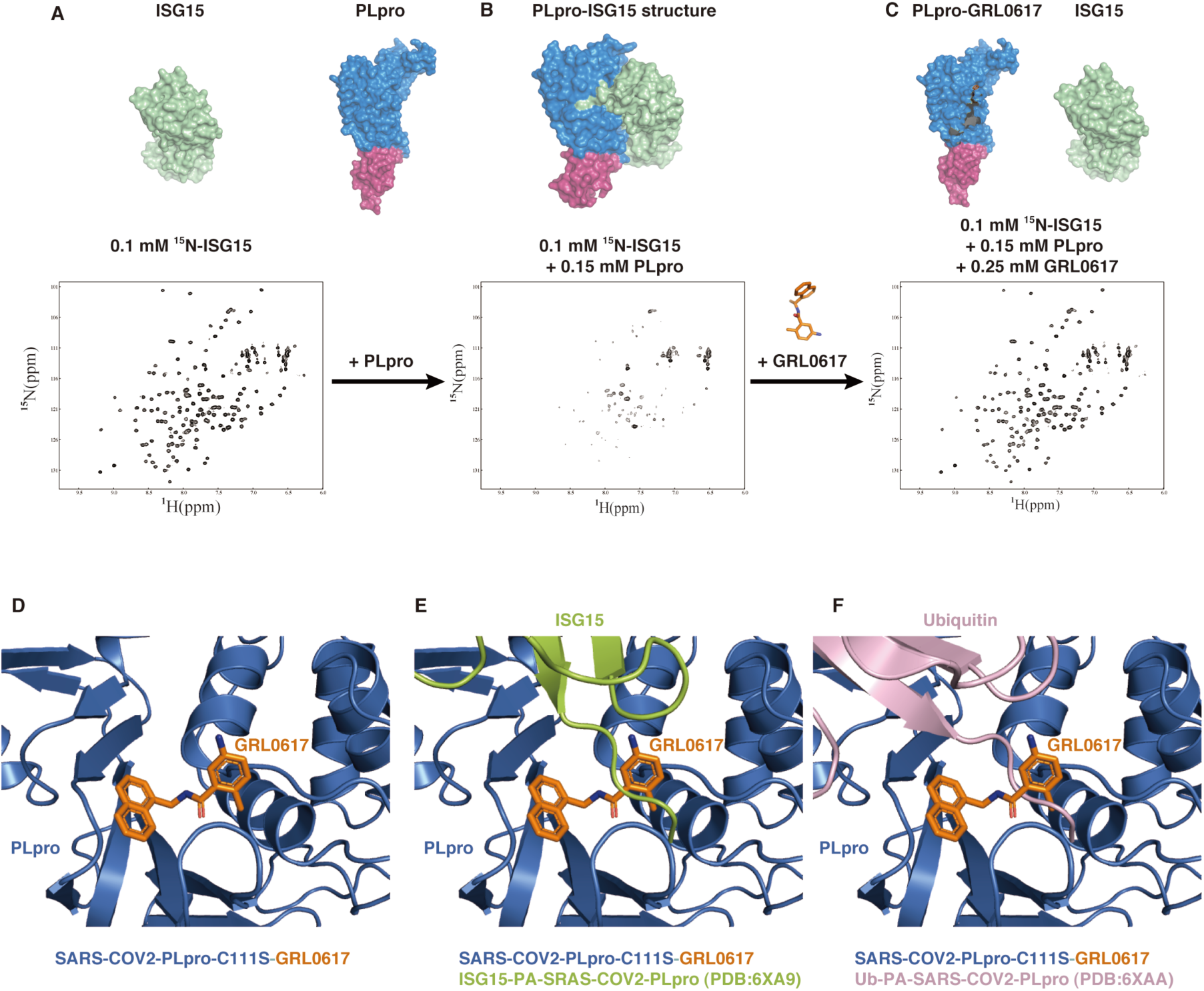
NMR studies show that GRL0617 blocks the binding of ISG15 to SARS-COV-2 PLpro. (A) ^1^H,^15^N-HSQC spectrum of ^15^N labeled ISG15; (B) HSQC spectrum of ^15^N labeled ISG15 (0.1 mM) and 0.15 mM PLpro. Peak broadening and peak intensity loss indicates binding of ISG15 to PLpro; (C) HSQC spectrum of ^15^N labeled ISG15 (0.1 mM) in the mixture of 0.15 mM PLpro and 0.25 mM GRL0617. Recovery of peak intensity suggests that GRL0617 binds to PLpro and displaces ISG15; (D) SARS-CoV-2 PLpro C111S/GRL0617 structure in cartoon model; (E) ISG15 in the complex structure of ISG15PA/SARS-CoV-2 PLpro (PDB 6XA9) was modelled on (D) by superposition, showing steric clash of GRL0617 with the ISG15 C-terminal tail; (F) Ub in the complex structure of UbPA/SARS-CoV-2 PLpro (PDB 6XAA) was modelled on (D) by superposition, showing steric clash of GRL0617 with the Ub C-terminal tail.

## Discussion

Our biochemical, biophysical and antiviral data of SARS-COV-2 PLpro support the claim that SARS PLpro is a promising drug target for COVID-19 treatment. PLpro’s multiple roles include cleaving polyprotein for virus maturation, dampening the host antiviral immune response by removing ISG15 modifications and dysregulating the host inflammation responses through cleaving K48-linked poly Ub chains from important proteins in viral invasion. Our study showed that the regulatory roles of PLpro can be inhibited by the small molecule GRL0617. Currently, drug discovery targeting the Mpro and the RNA-dependent RNA polymerase (RdRp) have brought some promising lead compounds to pharmaceutical development or even clinical trials, e.g. the RdRp inhibitor Remdesivir (*27, 28*). In our study, we show that PLpro is an equally promising target but more challenging because the co-crystal structure is hard to obtain and often irreproducible, like other coronaviral PLpros (*16*). Our co-crystal structure of PLpro C111S in complex with the potent inhibitor GRL0617 validated that SARS-COV-2 PLpro is a druggable target. Our structural characterization paves the way for future drug discovery targeting PLpro and provids a solid template for modeling and modifying potential inhibitors, including GRL0617 and its naphthalene analogs. Based on the structure, GRL0617 resides in the S3/S4 site of PLpro, naturally it will also inhibit the processing of viral polyproteins of SARS-CoV-2 since these viral polyproteins share the same substrate cleavage sequence with Ub and ISG15. Therefore, inhibition of SARS2-CoV-2 PLpro can simutaneouly prevent viral matureation and attaching host antiviral immune system. Our NMR study reveals that GRL0617 is a potent protein-protein interaction (PPI) inhibitor targeting PLpro. In comparison, Mpro is another highly explored drug target with several potent covalent inhibitors been reported (*11, 12, 14*). However, no covalent inhibitors of PLpro have ever been reported. Our study suggests that focusing on the discovery of non-covalent inhibitors could be an effective strategy targeting PLpro. Althrough the seven FDA-approved drug obtained in our screening show low potency against PLpro, we cannot rule out the potential of these drug to therapeutically treat COVID-19 because they may have higher antiviral activities by inhibiting other pathways in the virus lifecycle.

The structural rearrangement of the BL2 loop of PLpro upon binding GRL0617 provids a newly formed deeper pocket to better accommodate the compound. This binding induced pocket formation may imply that future virtual screening efforts should utilize the co-crystal structures. It is expected that this newly discovered pocket would become a hot spot for extensive efforts of drug optimization in the near future.

## Author Contributions

H.H. and G.Z. conceived the project. H.H., Z.F., J.T., Y.X. J.L. and Z.L. performed cloning, protein expression, isotope labeling and protein purification. Z.F. and H.H. conducted structure determination. J.T. and M.L. performed screening and enzyme assays with the help from Y.J.Z. B.H. and D.C. conducted cell-based assays as well as deubiquitination and deISGylation experiments. Y.Y. and H.Z. assisted in X-ray data collection and data analysis. J.T. performed NMR experiments with the help from Z.L. and X.N.. G.Z. and S.L. performed the antiviral and cytotoxicity assays. Y.J.Z. edited the manuscript. H.H. wrote the paper and everybody contributed to the writing.

## Acknowledgements

We thank Dr. Hans J. Vogel at the University of Calgary, Canada for critical reading and editing the manuscript. All NMR experiments were performed at the Beijing NMR Center and the NMR facility of National Center for Protein Sciences at Peking University. Research reported in this publication was supported in whole or in part by the National Natural Science Foundation of China (grant no: 21977009 to H.H.), the Shenzhen Science and Technology Innovation Committee (grant no: JCYJ20170412150913708 to H.H.), the Shenzhen Bay Laboratory Open Research Program (grant nos: SZBL2019062801010 to H.H. and SZBL202002271003 to G.Z.), the Guangdong Scientific and Technological Project (2020B1111340076 to G.Z.) and the Natural Science Foundation of the Guangdong Province (grant no: 2018A0303130091 to Z.F.).

