## Supplemental for "Structural basis for the inhibition of the papain-like protease of SARS-CoV-2 by small molecules"

Materials and Methods

Plasmids construction, protein expression, and purification

The protein sequence of SARS-CoV-2 PLpro_1563-1878_ of Nsp3 protein from SARS-CoV-2 (GeneBank: QHD43415) was codon optimized, synthesized and subcloned into pET28a vector with N-terminal His-tag and TEV protease site (General Biosystems, China). The C111S mutation was introduced into the PLpro with a QuikChange site-directed mutagenesis kit (Agilent Technologies, USA) and verified by sequencing. Human ISG15 was codon-optimized, synthesized and subcloned into pET28a with an N-terminal His-tag, SUMO-tag and TEV-protease site. Human Ub was codon-optimized, synthesized and subcloned into pET28a with an N-terminal His-tag and TEV-protease site.

The plasmids were subsequently transformed into *E. coli* BL21 (DE3) cells. Protein expression was carried out in LB medium. *E. coli* cells grow in LB medium at 37 °C until OD_600_ reaches 0.8-1.0. 0.5 mM Isopropyl β-D-1-thiogalactopyranoside (IPTG) and 100 mM ZnSO4 were added and cells keep growing overnight at 18 °C. For the production of ^15^N-labelled ISG15 and Ub, protein samples were prepared by growing in the bacteria in M9 medium containing ^15^NH4Cl.

Cell pellets were resuspended in buffer A (30 mM Tris, 400 mM NaCl, 30 mM Imidazole, 2 mM β-Me, pH 8.5) with the addition of 1 mM phenylmethylsulfonyl fluoride (PMSF) and lysed using sonication. Cell lysate was subsequently centrifuged at 18,000 rpm at 4 °C for 1h. The supernatant was further loaded on a Ni-NTA column and then purified using B (30 mM Tris, 400 mM NaCl, 300 mM Imidazole, 2 mM β-Me, pH 8.5) on an AKTA Pure purification system (GE Healthcare). A second step of purification was carried out on a Superdex 200 Gel filtration column using Buffer C (30 mM Tris, 100 mM NaCl, 1 mM DTT, pH 8.5, for crystallization) or Buffer D (30 mM Tris, 100 mM NaCl, 1 mM DTT, pH 7.4, for measuring the enzymatic activities and NMR tests). The fractions were pooled and concentrated to 10 mg/mL and stored at -80 °C.

*In vitro* deubiquitination assay

PLpro was diluted in dilution buffer (25 mM Tris-HCl (pH 7.5), 150 mM NaCl, 10 mM DTT, pH 7.5) and pre-incubated for 10 min at 23 °C. 1 μM of Ub chains with various linkages and lengths (Boston Biochem, USA) was prepared in reaction buffer (50 mM Tris-HCl, 50 mM NaCl, 5 mM DTT, pH 7.5). PLpro and Ub-chains were mixed by 1:1 ratio in the above reaction system and incubated at 37 °C for indicated period. Reactions were stopped by adding 2 × SDS-PAGE sample buffer and were subsequently subjected to SDS-PAGE and viewed by silver staining.

Immunoblotting for detection of ISGylation

HEK293T cells were co-transfected with plasmids encoding Myc-tagged ISG15, pcDNA3,1-UBE1L (E1), UBCH8 (E2), or 3XFlag-HERC5 (E3) for 24 h. Cell extracts were prepared in RIPA lysis buffer (20 mM Tris-HCl (pH 7.5), 150 mM NaCl, 1 mM EDTA, 1% NP-40, 1% SDS and protease inhibitor cocktail (Roche Diagnostics, Germany), pH 7.5. Protein concentrations were determined by Bradford assay (Bio-Rad Laboratories, CA, USA). Equal amount proteins (40 μg) were subjected to SDS-PAGE, followed by transferring to nitrocellulose membranes. Membranes were blocked with TBST containing 5% skim-milk for 1 h at room temperature and then overnight with ISG15 (Thermo Fisher Scientific), GFP, GAPDH (Abclonal, China), Ubiquitin (Cell Signal Technology, MA, USA) antibodies at 4 °C. After three washing steps, membranes were incubated with HRP-conjugated secondary antibodies (Santa Cruz Biotechnology, Dallas, TX) for 1 h at room temperature and visualized using ChemiDoc system (Bio-Rad Laboratories, CA, USA ).

PLpro activity assays using Peptide-AMC for HTS and IC_50_ determination

In this study, PLpro activity was monitored using the substrate peptide-AMC (Z-Arg-Leu-Arg-Gly-Gly-AMC, Cat. No. 4027158, Bachem Bioscience). Experiments were performed in 384-well black non-binding plates (Cat. No. 3575, Corning) with a final reaction volume of 50 μL. The assay buffer contained 50 mM HEPES, pH 7.4, 0.01% Triton X-100 (v/v), 0.1 mg/ml BSA, 2 mM DTT. PLpro was added to the plates at a final concentration of 100 nM. Enzyme reactions were initiated with 5 μL of peptide-AMC (final 50 μM) dissolved in the above assay buffer. Upon addition of peptide substrate, the fluorescence signals were monitored at 340 nm (excitation) and 450 nm (emission) with three-minute intervals in a 2104 EnVision Multilabel Plate Reader (PerkinElmer).

PLpro inhibitor screening using an FDA-approved drug library

An FDA-approved Drug Library (TargetMol, USA), including 2,111 compounds, was used. The 1^st^ round screening reaction mixture included 100 nM PLpro, 50 μM substrate and 100 μM compounds. The top 1.5% compounds of each plate were selected with a minimum 60% inhibition using the reaction in the DMSO as a control. Around 30 compounds were tested in the 2^nd^ round of screening using the same assay. 23 compounds were excluded due to their reactivities, insolubility or fluorescence interference. To determine the IC_50_ values of the remaining 7 compounds, a series of 8-point, 1:2 serial dilutions was performed from a highest starting concentration of 200 μM. Seven FDA drugs used in this study were purchased from TargetMol (USA). The data was fitted using GraphPad Prism.

Crystallization and data collection

The complex of GRL0617 (Cat. No. HY-117043, MCE) with PLpro C111S was crystallized by vapor diffusion in a sitting-drop format after a 20-h incubation of 9.5 mg/ml PLpro in the buffer (50 mM Tris, pH 8.5,100 mM NaCl) with 2 mM inhibitor at 4°C. Immediately before crystallization, the sample was clarified by centrifugation. A 0.75-μL volume of the enzyme-inhibitor solution was then mixed with an equal volume of well solution containing 5 mM Cobalt (II) chloride hexahydrate; 5 mM Cadmium chloride hemi (pentahydrate); 5 mM Magnesium chloride hexahydrate; 5 mM Nickel (II) chloride hexahydrate; 0.1 M HEPES, pH 7.5 and equilibrated against well solution at 12°C. Before data collection, crystals were soaked in a cryo solution containing well solution, 400 μM inhibitor, and 20% glycerol. Crystals were flash-frozen in liquid N2. All diffraction data were collected at an in-house light source Rigaku MicroMax-007 HF and indexed, integrated, and scaled using XDS(*1*).

Structure determination and data deposition

Structures were solved by molecular replacement using the apo-PLpro C111S structure (PDB 6WRH) as the template. The MOLREP(*2*) in CCP4(*3*) was used for MR. Structures were refined in PHINEX(*4*) with manual model building in Coot(*5*). Detailed statistics on the data collection and the final models for crystallographic analysis are shown in Table S1. Structural models were assessed using MolProbity(*6*). Structural alignments and graphical representations were generated using PyMOL(*7*). Coordinates and structure factors were deposited in the PDB under accession code 7CJM (GRL0617 bound PLpro).

Antiviral and cytotoxicity assay

1×10^4^ Vero E6 cells were seeded in triplicates in 96-well plates. After 20-24 hours, fresh medium with different concentrations (100, 50, 25, 12.5, 6.25, 3.125, 1.56 μM) of inhibitors, DMSO as a control, was replaced. For measuring the cytotoxicity, cells were incubated for 48 hours followed by the cell viability test with CCK8 reagent. For measuring the anti-viral activity, cells were kept in the medium with inhibitors for 1 h, followed by infection with SARS-CoV-2 virus strain BetaCoV/Shenzhen/SZTH-003/2020, which was clinically isolated from local patients, at MOI (multiplicity of infection) =0.01. After a 2-h incubation, the virus-compound mixture was subsequently removed, and fresh medium containing candidate compounds (100, 50, 25, 12.5, 6.25, 3.125, 1.56 μM) or DMSO was added, and cell growth was continued for 48 hours. The viral RNA was extracted from the supernatant medium and reverse transcription was performed to obtain transcripts. The linearized plasmid containing the S gene of the SARS-CoV-2 virus was transcribed *in vitro* and was used to prepare a standard curve to quantify the copy number of the virus. Primer and probe information: TaqMan primers for COVID-19 virus: 5'TCCTGGTGATTCTTCTTCAGG-3' and 5'-TCTGAGAGAGGGTCAAGTGC-3' and COVID-19 virus probe 5'-FAM-AGCTGCAGCACCAGCTGTCCA-BHQ1-3'. Data analysis was done with GraphPad Prism.

NMR Spectroscopy

NMR data were acquired at 25 °C on a 600-MHz Bruker AVANCE III spectrometer. The 600-MHz spectrometer was equipped with a 5-mm TCI CryoProbe. In NMR titrations, the samples of 0.1 mM ^15^N-labeled ISG15 were incubated in the presence or absence of 0.15 mM PLpro with or without the indicated concentration of GRL0617 (0.05 mM, 0.15 mM and 0.25 mM) were investigated in assay buffer containing 30 mM Tris, 100 mM NaCl, pH 7.4, 5% DMSO, 10% D_2_O. For Ub titrations, the samples of 0.1 mM ^15^N-labeled Ub were incubated with PLpro (0.1 mM, 0.2 mM and 0.3 mM). ^1^H,^15^N-HSQC titration spectra were collected for all the samples. All of the NMR spectra were processed using NMRPipe/NMRDraw and further analyzed using NMRView.

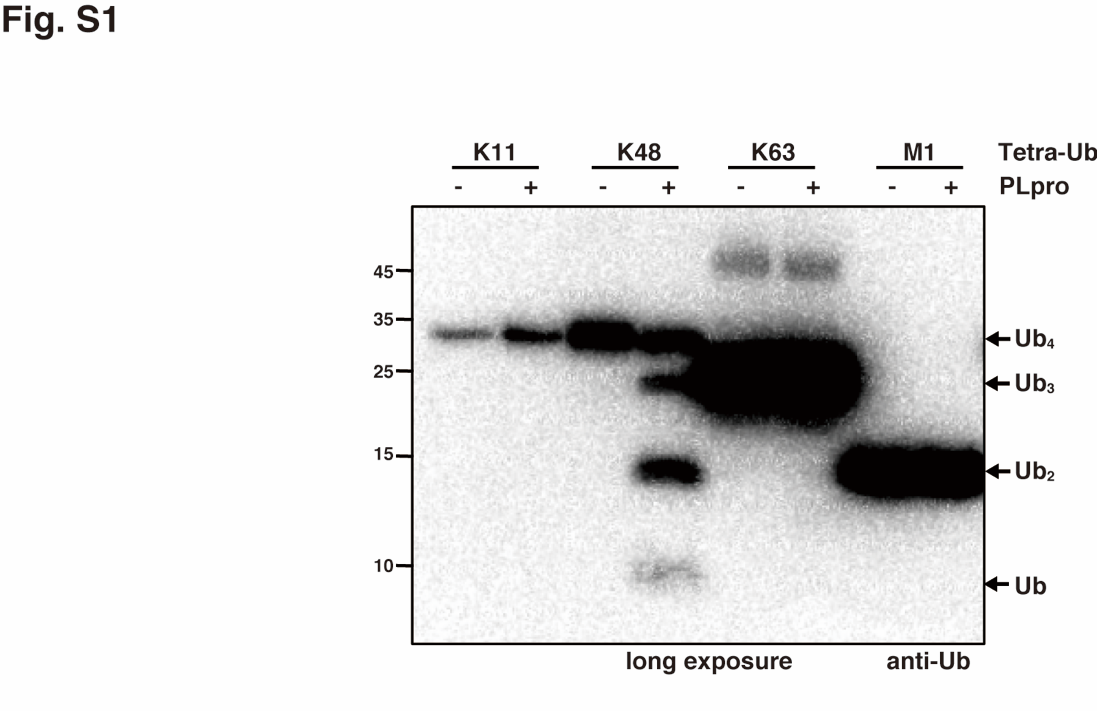

**Fig. S1 Linkage preference of SARS-COV-2 PLpro enzyme activity with 2 sec exposure.** PLpro was incubated with K11-, K48-, K63-tetra-Ub and M1-di-Ub for 30 min at 37℃, then analyzed by SDS-PAGE and immunoblotting with 2 s exposure.

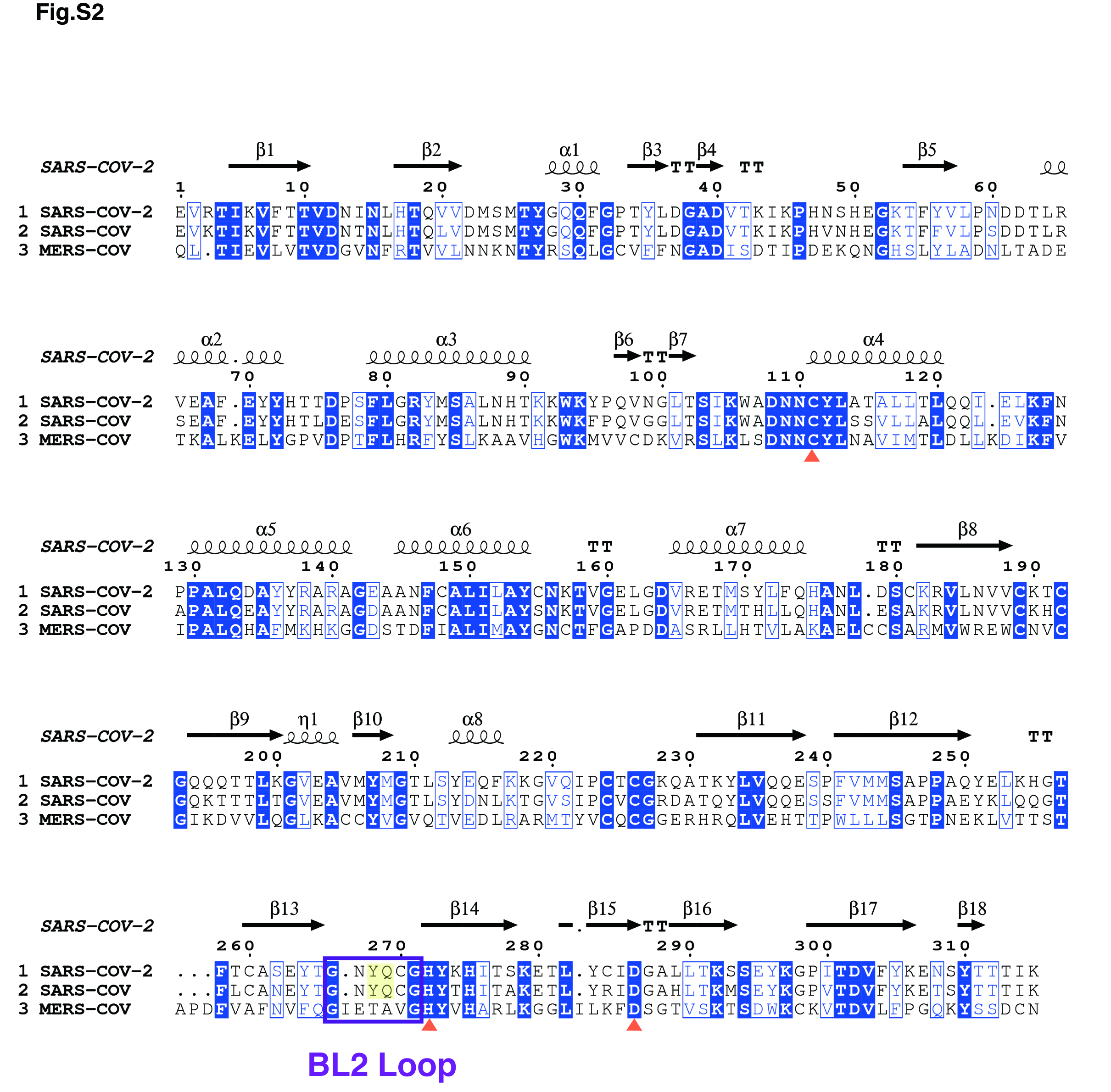

**Fig. S2 Sequence alignment of SARS-CoV-2/SARS/MERS PLpro.**

Sequence alignment generated with MUSCLE/ESPRIPT(*8, 9*) aligning PLpro sequences from SARS-CoV-2, SARS, MERS. Sequence numbering and secondary structure elements are shown according to the apo-structure of SARS-CoV-2 PLpro C111S (PDB ID: 6W9C chain A). The catalytic triad residues are labeled with red trigonometry; the BL2 loop is shown with purple frame; two residues on the BL2 loop which has structural rearrangement upon GRL0617 binding are colored in yellow. α = α-helix, β = β-strand.

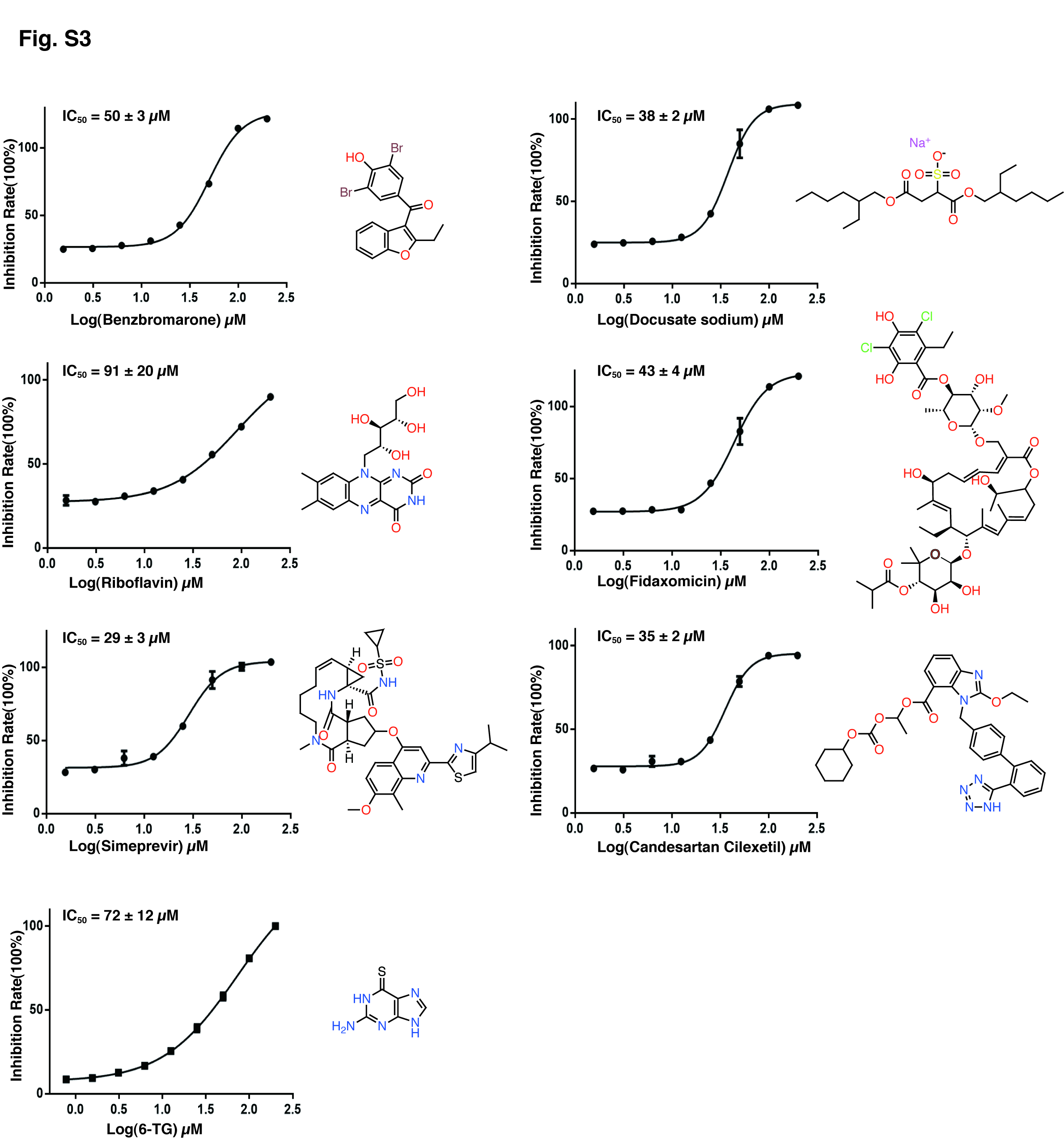

**Fig. S3 The inhibitory activity of FDA-approved drugs against PLpro of SARS-CoV-2.** Dose-response inhibition profile of drugs for half-maximum inhibitory concentration (IC_50_) values were fitted using Graphpad prism. Data are presented as the means ± SEM, n = 3.

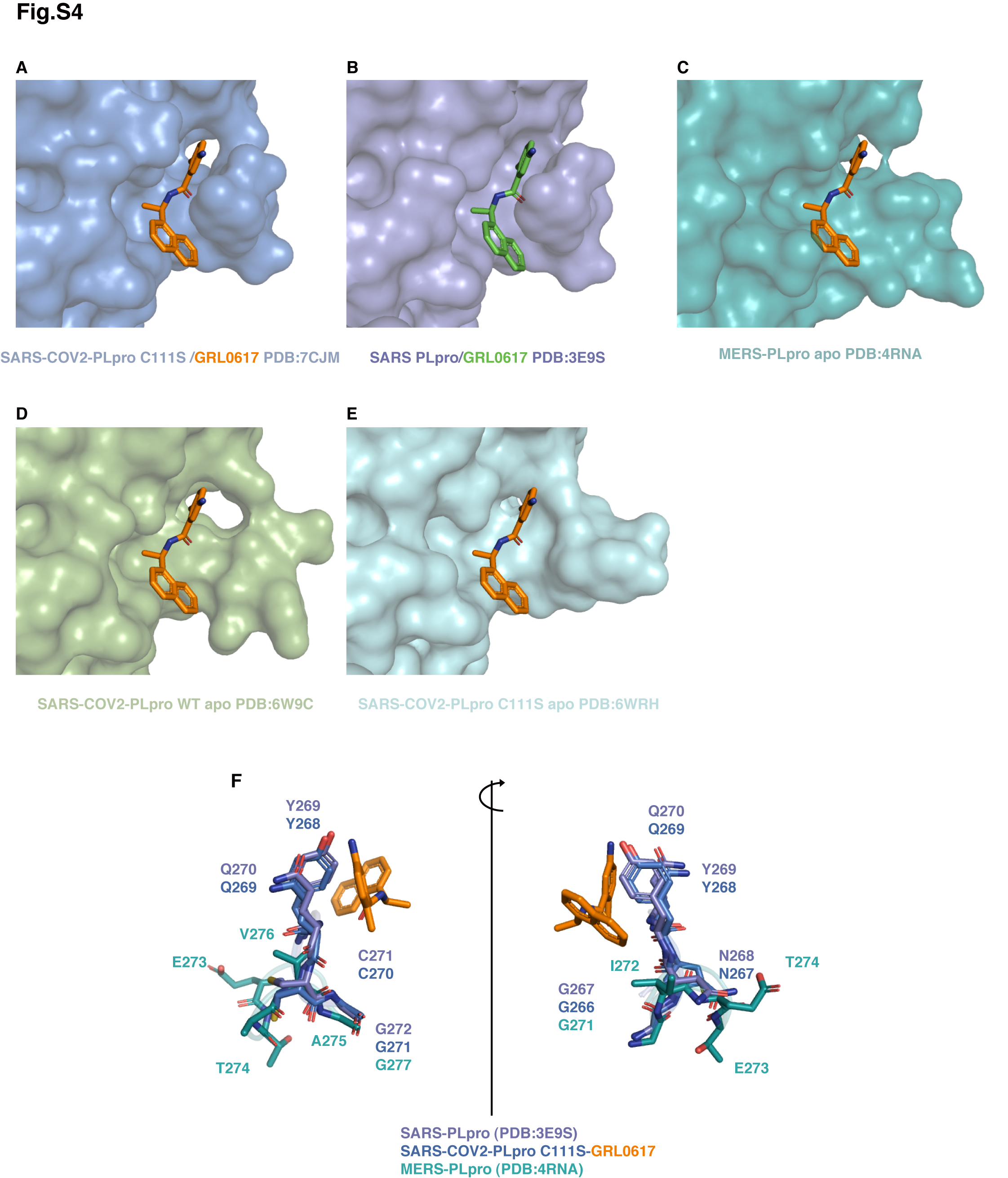

**Fig. S4 Surface representations of the GRL0617-Binding site of PLpro and the corresponding region of known viral PLpro crystal structures.**

(A) SARS-CoV-2 PLpro C111S/GRL0617; (B) SARS-CoV PLpro in complex with GRL0617 (PDB ID: 3E9S); (C) MERS PLpro superposition with (A) (PDB ID: 4RNA); (D) apo-SARS-CoV-2 (PDB ID: 6W9C) superposition with (A); (E) apo-SARS-CoV-2 PLpro C111S (PDB ID: 6WRH) superposition with (A) (F) Superpositions of BL2-Loop of SARS-COV-2/SARS-CoV/MERS PLpro. BL2 loop in MERS PLpro is not as conserved as in SARS-COV2 and SARS.

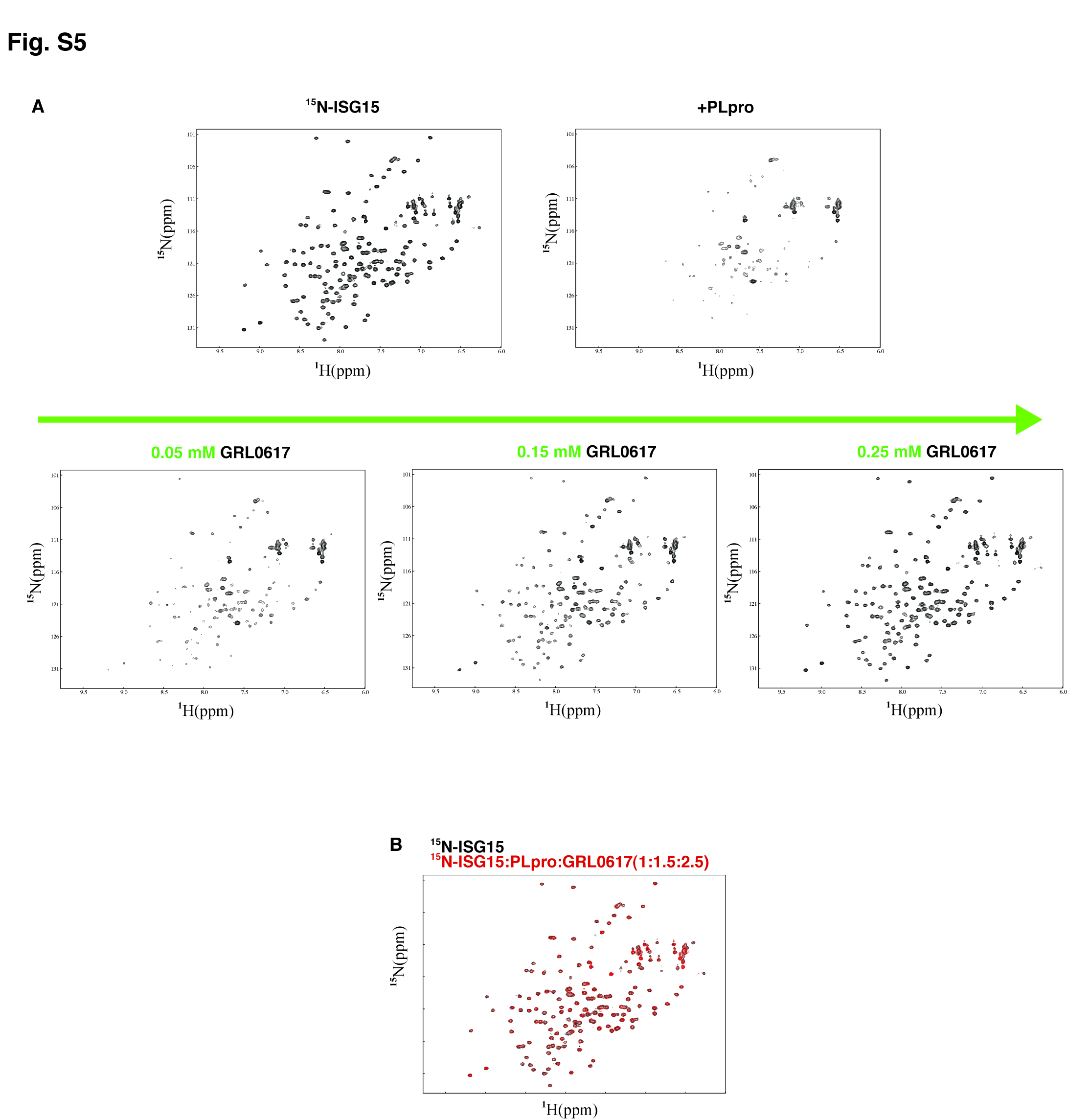

**Fig. S5 GRL0617 blocks the binding of ISG15 to PLpro in a dose dependent manner.** (A) Addition of PLpro caused peak disappearing of ^15^N-ISG15 due to binding. With the increase concentrations of GRL0617, the cross-peaks of ^15^N-ISG15 gradually reappeared. (B) Superposition of ^1^H, ^15^N-HSQC spectra at the indicated molar ratios for ^15^N-ISG15 (black) versus ^15^N-ISG15 : PLpro : GRL0617 (red).

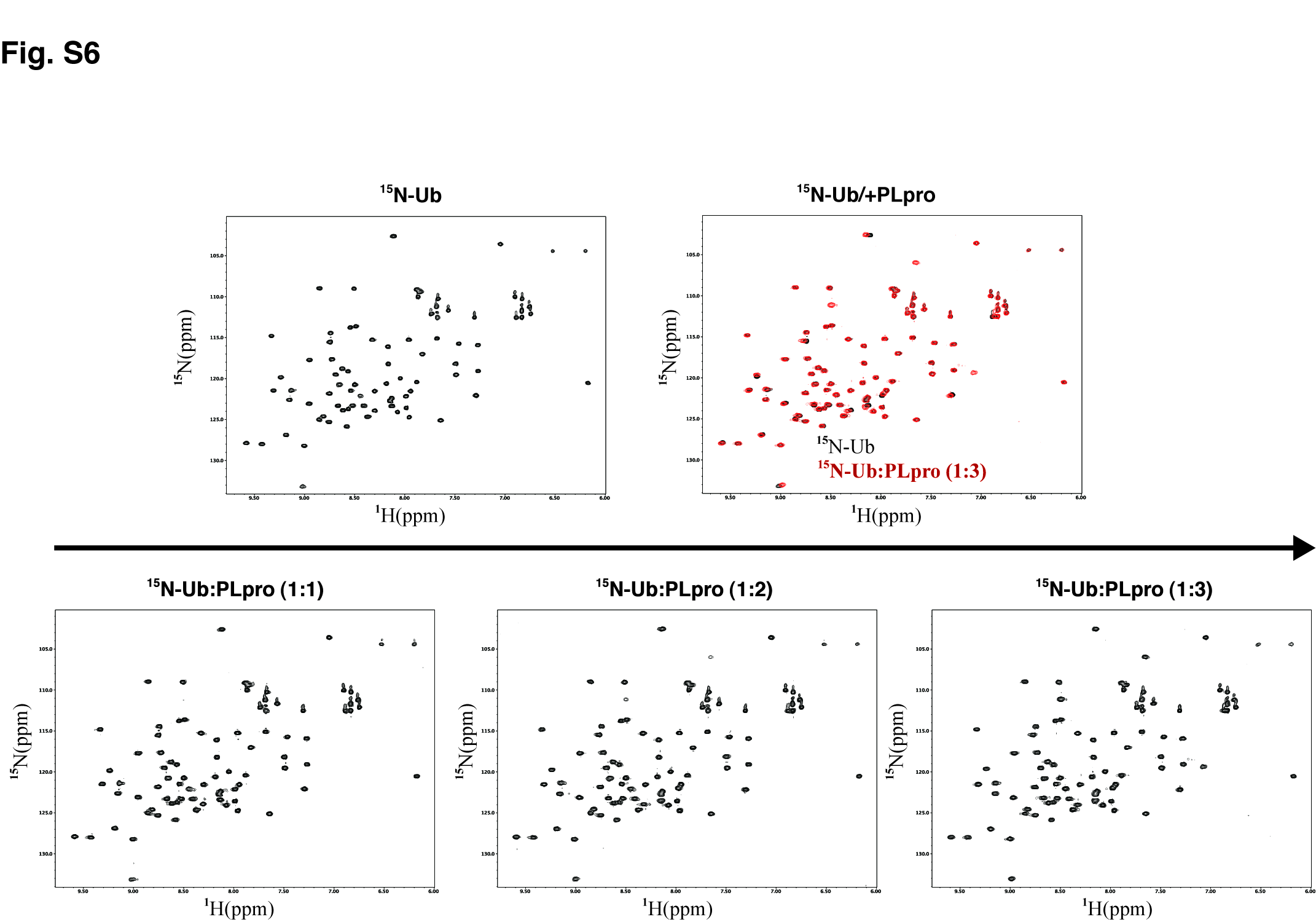

**Fig. S6 ^1^H, ^15^N-HSQC spectra at the indicated molar ratios for ^15^N-ISG15 (black) in mixture with SARS-CoV-2 PLpro.** Marginal peak shifting at 1/3 molar ratio suggests very weak binding between monoUb and PLpro under *in vitro* conditions.

**Table S1 Crystallographic data for the co-crystal structure of SARS-CoV-2 PLpro in complex with GRL0617.**

| **Protein/Ligand** | **Plpro+GRL0617** |
| --- | --- |
| **PDB entry** | 7CJM |
| **Data Collection** |  |
| Space group | I 41 2 2 |
| Cell dimentions |  |
| a,b,c(Å) | 112.29, 112.29, 220.38 |
| α,β,γ(°) | 90 90 90 |
| Resolution range(Å) | 22.14  - 3.2 (3.314  - 3.2) |
| Rmerge(%) | 19.0 (48.7) |
| I/sigma(I) | 12.92 (4.27) |
| Completeness (%) | 99.05 (99.83) |
| Redundancy | 6.3 (6.6) |
| **Refinement** |  |
| Resolution(Å) | 3.2 |
| Number of Unique reflections used for refiement | 11920 (1178) |
| Rwork/Rfree(%) | 26.78/28.62 |
| No. of non-hydrogen atoms | 2533 |
| Macromolecules | 2507 |
| Protein residues | 316 |
| Ligand | 26 |
| RMSDs |  |
| Bond lengths(Å) | 0.006 |
| Bond angles(°) | 0.86 |
| Ramachandran plot(%) |  |
| Favored | 95.54 |
| Allowed | 4.46 |
| Outliers | 0 |
